# A method for estimating the response of nursery-grown Atlantic Forest tree seedlings to water deficit

**DOI:** 10.64898/2026.07.07.737083

**Authors:** Larissa Cerqueira Dias Rodrigues, José Antônio Pimenta, Fátima Arcanjo, Alba Lúcia Cavalheiro, Halley Caixeta de Oliveira, José Marcelo Domingues Torezan

## Abstract

Global climate change has increased the frequency and intensity of drought events, making it urgent to understand how native species respond to water deficit (WD). In biodiverse environments such as tropical forests, simple methods are needed to study multiple species simultaneously. This can help predict how natural environments will respond to climate change and guide the strategic selection of drought-resistant species for reforestation. This study aimed to: (1) adapt an existing simple and inexpensive method to apply a controlled WD on tree seedlings from tropical species commonly produced in nurseries for restoration projects, suitable for greenhouse experiments; and (2) evaluate the effectiveness of this method in generating ecophysiological responses to WD that allow the estimation of species’ drought resistance. Ten native tree species from the Semideciduous Seasonal Forest (SSF), a phytophysiognomy of the Atlantic Forest, were selected. An existing method was adapted to implement capillary irrigation, in which the bases of the seedling tubes were placed in floral foam blocks positioned inside 15 L plastic containers filled with water. A gradual and severe WD was applied to five seedlings of each species by removing all water from the containers, leaving only the water retained in the saturated floral foam available for plant uptake. The remaining seedlings were maintained well-watered (containers full and foam saturated) as the control group. Stomatal conductance (*g*_s_) was measured daily for all seedlings until they reached 50% or less of their initial *g*_s_ (i*g*_s_); at this point, stem water potential (Ψw) was measured. Both *g*_s_ and Ψw differed significantly among treatments and species (p < 0.01). *Ficus guaranitica* and *Heliocarpus popayanensis* were the only species that did not show significant Ψw differences between treatments, indicating higher drought resistance. In contrast, *Campomanesia xanthocarpa* and *Eugenia uniflora* had the lowest Ψw values under WD, suggesting lower drought resistance. The remaining species were distributed along a gradient of responses to WD. Additionally, no correlation was found between Ψw and g_s_ at 50% i*g*_s_ in the WD group (*rho* = 0.16, p = 0.26). The method proved effective in inducing controlled WD and generating measurable ecophysiological responses, offering a useful tool for screening native species for drought resistance.

## 1 INTRODUCTION

Drought events and intense heat waves are expected to increase in intensity and frequency around the globe due to climate change (IPCC, 2023). In South America, the major aggravating factor is deforestation, mostly related to agricultural production for exportation (Salazar et al., 2016). Those changes in land use exacerbate the impacts of climatic changes and make the region one fundamental actor in the future of the world’s economy and food production (IPCC, 2022). International emission reduction programs have primarily focused on the detection and monitoring of deforestation (Pearson et al., 2017). Reducing carbon emissions from deforestation and forest degradation is a key strategy for mitigating climate change (Bustamante et al., 2019). However, in Brazil, much of the native vegetation has already been altered or lost, particularly in the Atlantic Forest, where only about 12.4% of mature and well-preserved forest cover remains (SOSMA, 2024).

As a result, ecosystem restoration has increasingly been adopted as a strategy for climate change mitigation through CO_2_ sequestration (Koch and Kaplan, 2022). Active restoration, which involves the planting of seedlings, is especially important in these highly fragmented landscapes (Holl, 2023). However, an ongoing challenge, particularly in biodiverse tropical regions, is the selection of native species that are resilient to increasing climatic extremes, especially severe droughts predicted under future climate scenarios. Identifying drought-resistant species is therefore essential to improve the success and long-term sustainability of restoration initiatives.

The ecological resistance to drought can be related to some physiological responses of plants, such as: (i) low stomatal conductance (*g*_s_), associated with conserving the available soil water by reducing leaf transpiration rates (Kazmierczak et al., 2015); and (ii) higher leaf water potential (Ψw)— that is, greater water availability— when *g*_s_ is reduced by 50%, which may be related to the prevention of functional loss and xylem damage (Brodribb and Holbrook, 2003). In this context, a widely recognized framework proposed by McDowell et al. (2008) distinguishes two key physiological mechanisms underlying plant mortality under drought: the isohydric and anisohydric strategies. Isohydric species tend to close their stomata early during drought, which limits water loss but increases their vulnerability to carbon starvation. In contrast, anisohydric species keep their stomata open for longer, even under low water potentials, thereby maintaining carbon assimilation but increasing the risk of hydraulic failure.

Nonetheless, studies assessing drought resistance across several tropical tree species are still scarce. This gap is partly due to the high cost and logistical complexity of simulating water deficit (WD) in greenhouse conditions (Earl, 2003), as well as the technical challenges involved in measuring certain physiological responses that are not easily applicable to a large number of species. While some studies have proposed simplified approaches to estimate drought tolerance (Fernández and Reynolds, 2000, Marchin et al., 2020), these often rely on large plants grown in pots of up to 6 liters, requiring seedling transplantation and long cultivation periods.

Alternatively, studies using smaller plants typically rely on seedlings grown specifically for experiments (Tiepo et al., 2018, Calzavara et al., 2021, Do Carmo et al., 2021), which involves seed collection, germination, and transplanting into larger containers, with daily manual watering for several months. This process is labor-intensive, time-consuming, and prone to high seedling mortality. Moreover, manual irrigation makes it difficult to standardize soil moisture levels, potentially compromising the consistency and reliability of the results.

In short, a major limitation of previous approaches is the difficulty of scaling up experiments in a short timeframe. Therefore, it is essential to develop simple and inexpensive methodologies that can be applied on a large scale to accelerate the generation of the knowledge needed to interpret species-rich environments such as tropical forests. Accordingly, this study aimed to 1) adapt an existing simple and inexpensive method to apply a controlled WD on tree seedlings from tropical species commonly produced in nurseries for restoration projects, suitable for greenhouse experiments; and 2) evaluate the effectiveness of this method in generating common ecophysiological responses to WD that allow the estimation of species’ drought resistance. To achieve this, seedlings of ten tree species native to the Semideciduous Seasonal Forest, a formation within the Atlantic Forest, were selected from a local nursery.

## 2 MATERIAL AND METHODS

### 2.1 Plant material

Two experiments were conducted between December 2022 and January 2023, at a greenhouse in the Center of Biological Sciences at Londrina State University. Ten tree species, all native to the Seasonal Semidecidual Forest, the hinterland form of the Brazilian Atlantic Forest, and commonly used in restoration projects were selected. To maximize phylogenetic diversity and increase the chance of detecting different levels of responses to WD that may be related to drought resistance, the species were chosen from 10 different genera and eight families (Table 1). Five species were used in each experiment and for each species, 10 seedlings were selected; five were kept well-watered (WW) (Control) and five were subjected to WD.

**Tabel 1.** List of native species of the Semideciduous Seasonal Forest, used in experiments with severe water deficit with seedlings produced in nurseries.

| Species | Code | Family |
| --- | --- | --- |
| *Aegiphila integrifolia* (Jacq.) Moldenke | AEI | Lamiaceae |
| *Allophylus edulis* (A.St.-Hil. et al.) Hieron. ex Niederl. | ALE | Sapindaceae |
| *Aspidosperma polyneuron* Müll. Arg. | ASP | Apocynaceae |
| *Campomanesia xanthocarpa* (Mart.) O.Berg | CAX | Myrtaceae |
| *Eugenia uniflora* L. | EUU | Myrtaceae |
| *Ficus guaranitica* Chodat | FIG | Moraceae |
| *Gallesia integrifolia* (Spreng.) Harms | GAI | Phytolaccaceae |
| *Heliocarpus popayanensis* Kunth | HEP | Malvaceae |
| *Maclura tinctoria* (L.) D.Don ex Steud. | MAT | Moraceae |
| *Schinus terebinthifolia* Raddi | SCT | Anacardiaceae |

The seedlings were produced in plastic tubes with an internal volume of 50 ml, whose available water content (6-10 ml, depending on substrate) can run out in a few hours. The substrate used is formulated with organic compost made from plant remains, vermiculite, and encapsulated fertilizer (Osmucote®). This type of seedling production is commonly used throughout the Atlantic Forest region.

All the seedlings used were at the end of the growth period, maintained under 50% shade screen in the nursery, just before entering for acclimatization in the full sun. We selected seedlings of similar canopy size, exposed roots and avoiding tubes with gaps in the substratum.

### 2.2 Experimental design

We adapted the method of capillarity irrigation system described by Marchin et al. (2020) which is itself an adaptation of the method first described by Snow and Tingey (1985) and already modified by Fernández and Reynolds (2000). Here, we used “floral foam”, which is a phenolic, non-elastic foam, commonly used for Ikebana and other flower arrangements, in the form of blocks measuring 22 cm x 10 cm x 7 cm. Foam blocks were placed in 15 L containers filled with water; three foam blocks were arranged in each container. After that, three seedlings were allocated in each foam (Fig. 1).

**Fig. 1.**
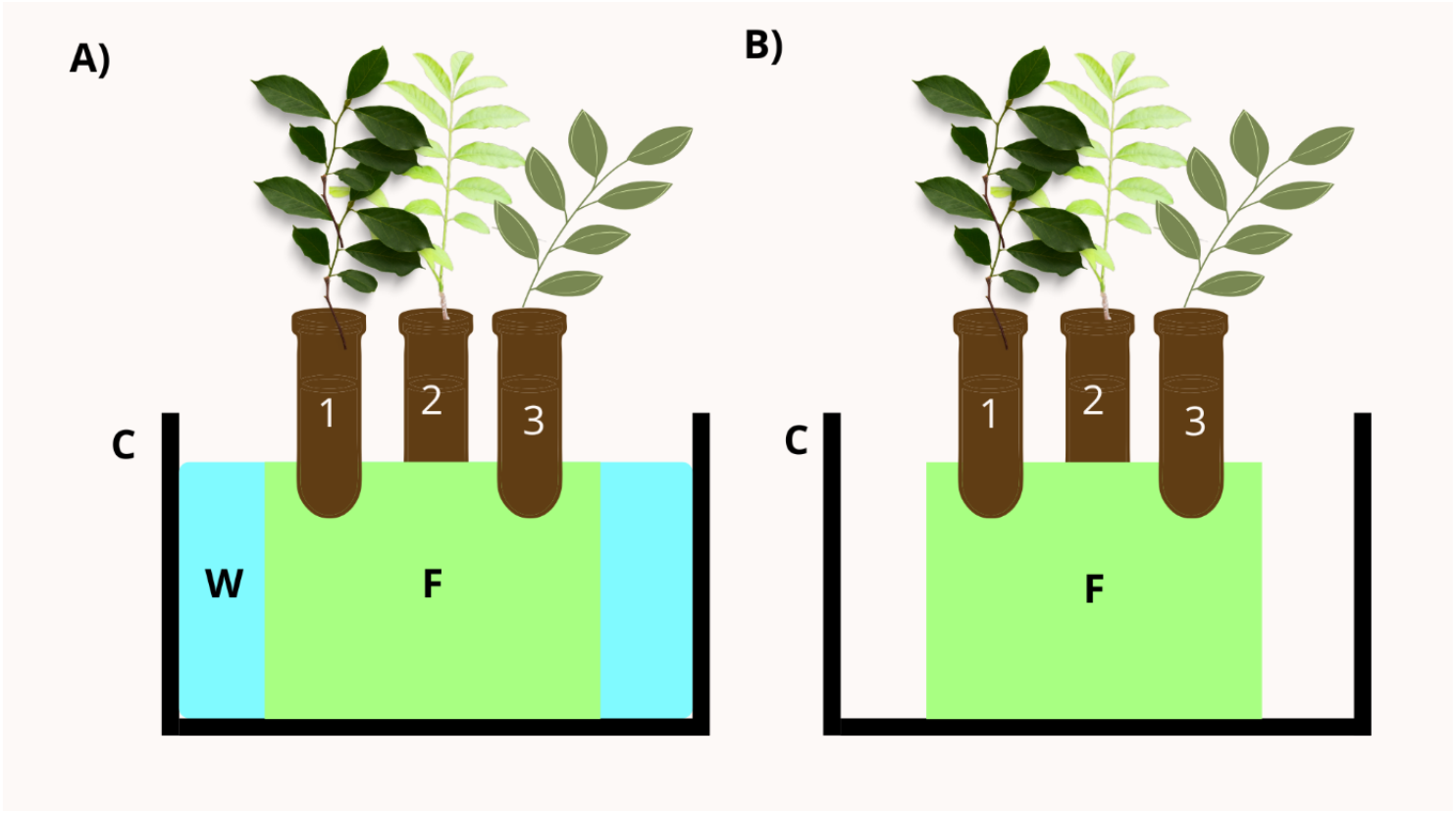
Diagram of the simple water deficit method, adapted from Marchin et al., 2020 (A Simple Method for Simulating Drought Effects on Plants). **A**) Control container-capillary irrigation is used to control soil water content of nursery seedlings (1,2,3), which are placed above a solid column permeable to water. **B**) Container in severe water deficit, all the water in the container was removed, leaving only the water from the soaked floral foam for seedlings to use. (F) commercial floral foam inside a plastic container (C) filled with water (W).

It is important to note that when choosing the material, two main points must be observed: (1) there is an adequate contact area between the soil at the bottom of the tube and the surface of the foam and (2) the pore size of the floral foam is suitable to transport water to the desired height by capillarity (Supplementary Material Fig.S1).

To ensure uniform spacing and prevent canopy overlap, three interspersed wells were made in each floral foam blocks using empty tubes (Supplementary Material Fig.S2). Seedlings from different species were randomly assigned to the wells. All containers underwent a seven-day acclimatization period in the greenhouse with the water table just below the foam block upper side and replenished every two days to keep the substrate WW. This approach reduced labor by eliminating the need for frequent manual watering.

According to Marchin et al., (2020), this method is able to maintain the volumetric soil water content between 30 and 39% in the Control tubes, which is considered field capacity (∼35%). Thus, at the beginning of the experiment, all seedlings (WD and Control) were at field capacity, and baseline physiological measurements were taken. Then, half of the containers were maintained with the water table just below the top of the foam block, while in the other half, the water table was reduced to zero, initiating the WD treatment. Although the imposed WD was severe, it progressed gradually, as the seedlings continued to extract water from the foam, which was slowly depleted over time.

### 2.3 Plant physiological measurements

Stomatal conductance was measured daily between 7:30 a.m. and 10:30 a.m., following the random position of the seedlings in the containers. Measurements were taken on the youngest fully expanded and undamaged leaf of each plant using a porometer (SC-1 leaf porometer, Decagon Devices, Pullman, WA, EUA). The experiment was concluded when seedlings in the WD treatment reached a g_s_ value less than or equal to 50% of their initial g_s_ (*ig*_s_). Once they reached this threshold, stem water potential (Ψw) was measured using a pressure chamber (SAPS II, model 3115, Saint Barbara, CA, EUA).

### 2.4 Statistical analysis

To analyze physiological responses to WD, two generalized linear models (GLMs) were fitted using the Gamma distribution with an inverse link function (Zuur et al., 2009). In the first model, *g*_*s*_ measured on the third day of the experiments − the last day with complete *g*_*s*_ data for all individuals − was modeled as the response variable to evaluate early stomatal regulation, which can be an indicative of drought resistance. The fixed effects included species and treatments (WW or WD).

In the second model to assess interspecific variation in Ψw with a loss greater than or equal to 50% of *ig*_s_, Ψw was modeled as the response variable and species and treatments as explanatory variables. To satisfy the assumptions of the Gamma distribution, Ψw values were log-transformed using the formula *log* (Ψw + 5), to ensure all values were positive before transformation. Models were fitted using the “glm.nb” function from the *MASS* package (Venables and Ripley, 2002). Post-hoc pairwise comparisons were conducted using the “emmeans” and “cld” functions from the *emmeans* (Russell, 2019) and *multcomp* (Hothorn, Bretz and Westfall, 2008) packages, with significance set at p < 0.05.

Generalized linear mixed models (GLMMs; Zuur et al., 2009) were also tested for each of the response variables mentioned above, using the Gamma distribution family and considering species and treatments as an explanatory factor. In the GLMMs, the experiment was included as a random variable to account for natural climatic variability between the two experiments. The best model was selected based on the lowest values of the Akaike Information Criterion (AIC) (Burnham and Anderson, 2002), the Bayesian Information Criterion (BIC) (Schwarz, 1978), and maximum likelihood (Supplementary Material Table S1). The GLMs showed better fit to the data as the random effect did not have significant influence on the models. Therefore, the results of the GLMs will be presented.

Additionally, Spearman’s rank correlations were performed to investigate the relationship between the final *g*_s_ measurement (after a ≥ 50% reduction from the i*g*_s_) and stem Ψw at the same *g*_s_ threshold. Correlations were conducted separately for individuals of all species within each treatment and were considered significant at p < 0.05. The strength of the correlations was interpreted according to Callegari-Jacques (2003): no correlation (*rho* = 0), weak (0 < *rho* ≤ 0.3), moderate (0.3 < *rho* ≤ 0.6), strong (0.6 < *rho* ≤ 0.9), and very strong (0.9 < *rho* ≤ 1). All statistical analyses were performed in software R (4.4.0). All the graphs were created with the *ggplot2* package (Wickham, 2009). The summarized data used in the analyses are shown in Supplementary Material Table S2.

### 3 RESULTS

The GLM analysis of *g*_s_ data collected on the third day of the experiment revealed significant differences between treatments and among species, as well as a significant interaction between treatments and species (Table 2, Supplementary Material Table S3).

**Table 2.** GLM results comparing the stomatal conductance (*g*_s_) of ten native species of the Semideciduous Seasonal Atlantic Forest on the third day of severe water deficit experiment in a greenhouse. Five individuals of each species were submitted to a water deficit treatment and the other five continued to be irrigated by capillarity (control group). Df: degrees of freedom; Deviance: model deviance; Residual Df: residual degrees of freedom; Residual Deviance: residual deviance; Pr(>Chi): p-value associated with the chi-squared statistic.

| Response Variable | Explanatory Variables | Family | Df. | Deviance | Resid. Df | Resid. deviance | Pr(>Chi) |
| --- | --- | --- | --- | --- | --- | --- | --- |
| $g_s$ | Species | Gamma | 9 | 14.90 | 89 | 28.91 | < 0.01 |
|  | Treatments |  | 1 | 8.83 | 88 | 20.08 | < 0.01 |
|  | Species:treatments |  | 9 | 3.52 | 79 | 16.55 | 0.03 |

Multiple pairwise comparisons of the *g*_s_ data revealed the formation of distinct groups, highlighting interspecific variation in responses to WD (Supplementary Material Table S4). *C. xanthocarpa, M. tinctoria*, and *A. edulis* showed the lowest *g*_s_ under WD on the third day of the experiments. Among them, only *M. tinctoria* exhibited a significant difference between treatments (Fig.2, Supplementary Material Table S5).

*C. xanthocarpa* and *A. polyneuron* stood out as the species with the lowest *g*_*s*_ even under WW conditions. In contrast, *H. popayanensis, F. guaranitica*, and *S. terebinthifolius* had the highest *g*_s_ under WD on day three, with only *S. terebinthifolius* showing a significant difference between treatments. Although *F. guaranitica* did not exhibit a statistically significant difference, its *g*_s_ dropped by more than 50% compared to its initial value. Thus, *H. popayanensis* emerges as the only species that maintained high *g*_s_ without showing any significant treatment effect on the third day of experiment. The remaining species were distributed along a spectrum of responses to WD (Fig. 2).

**Table 3.**
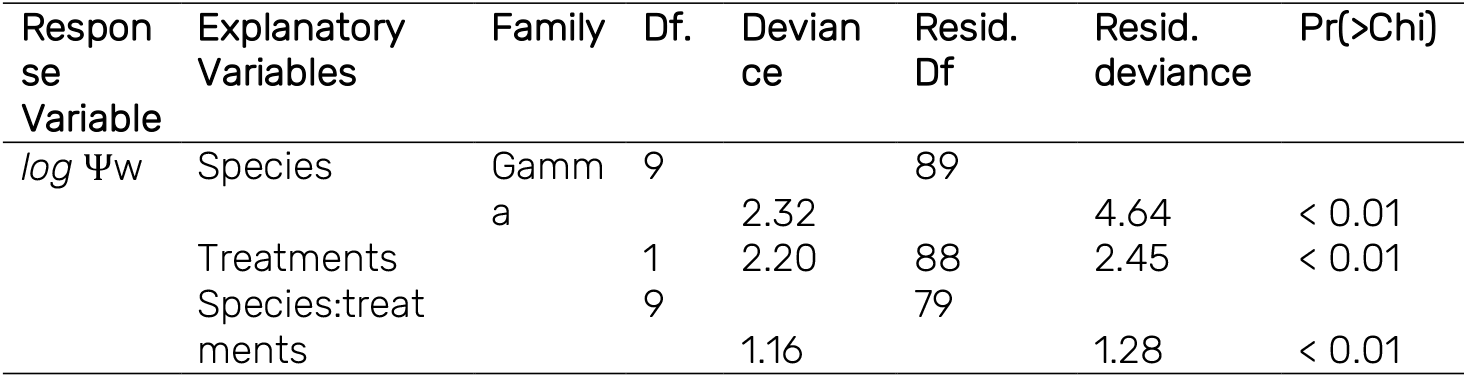
GLM results comparing the water potential (Ψw) of ten native species of the Semideciduous Seasonal Atlantic Forest measured when stomatal conductance had decreased by ≥50% from its initial value in a greenhouse experiment. Five individuals of each species were submitted to a water deficit treatment and the other five continued to be irrigated by capillarity (control group). Df: degrees of freedom; Deviance: model deviance; Residual Df: residual degrees of freedom; Pr(>Chi): p-value associated with the chi-squared statistic

**Fig. 2.**
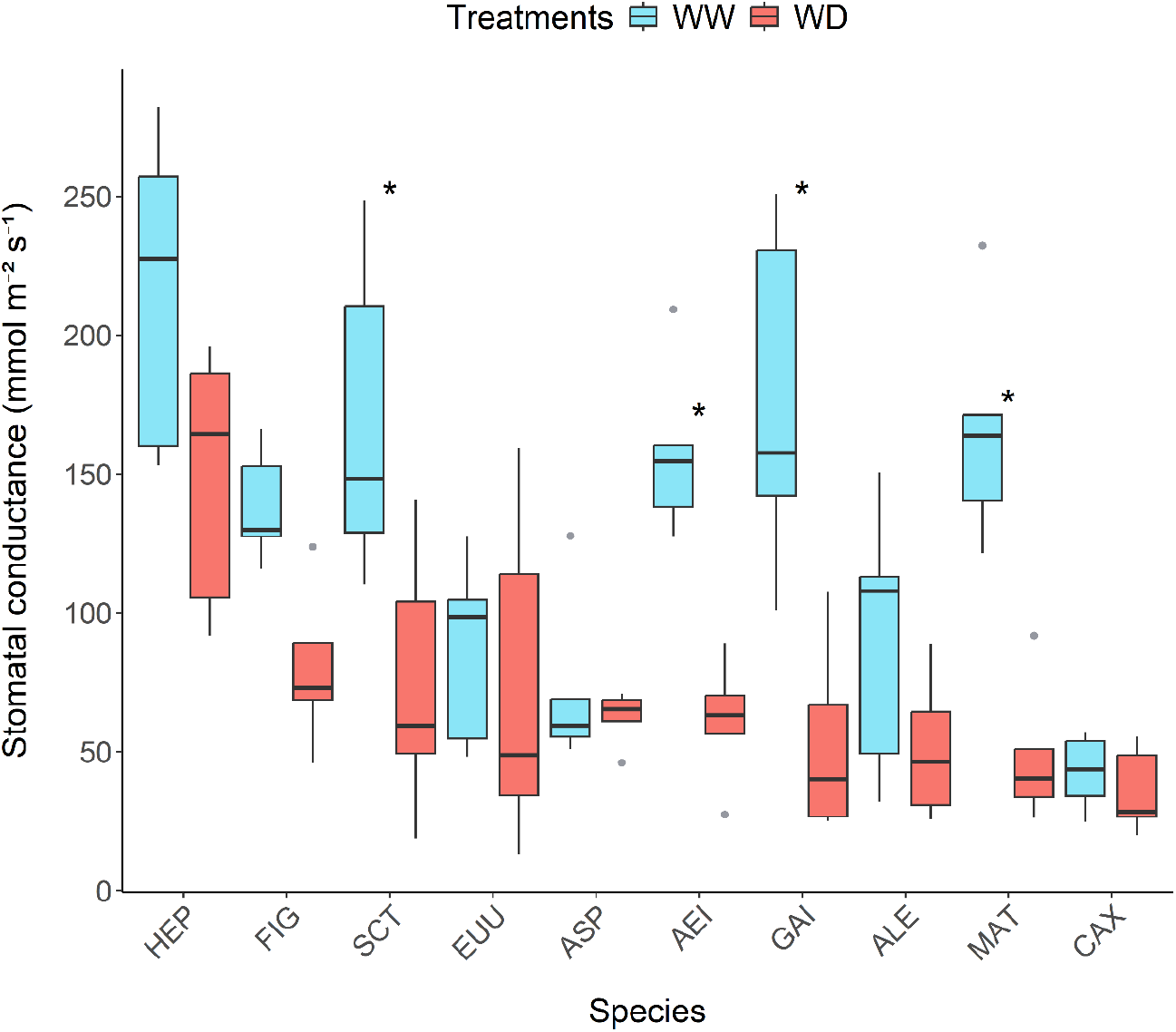
Stomatal conductance (g_s_) of seedlings from ten native species of the Semideciduous Seasonal Forest under well-watered (WW-blue bars) and water-deficit (WD-red bars) conditions on the third day of a severe WD experiment conducted in a greenhouse. AEI-*Aegiphila integrifolia* (Jacq.) Moldenke, ALE-*Allophylus edulis* (A.St.-Hil. *et al*.) Hieron. ex Niederl., ASP-*Aspidosperma polyneuron* Müll. Arg., CAX-*Campomanesia xanthocarpa* (Mart.) O.Berg. EUU-*Eugenia uniflora* L., FIG-*Ficus guaranitica* Chodat, GAI -*Gallesia integrifolia* (Spreng.) Harms, HEP-*Heliocarpus popayanensis* Kunth, MAT-*Maclura tinctoria* (L.) D.Don ex Steud., SCT-*Schinus terebinthifolius* Raddi. ^*^ Indicates significant differences between treatments.

The GLM results for Ψw data also revealed significant main effects of treatments and species, and a significant interaction effect (Tabel 3, Supplementary Material Table S6), reinforcing the results found about *g*_s_.

Multiple pairwise comparisons between species using the Ψw data also revealed the formation of several distinct groups, confirming a wide range of responses to WD (Supplementary Material Table S7). The species *E. uniflora* and *C. xanthocarpa* showed the lowest water potential (Ψw) under water deficit (WD), while *H. popayanensis* and *F. guaranitica* had the highest values and were the only species that maintained similar Ψw between treatments (Fig. 3, Supplementary Material Table S8).

**Fig. 3.**
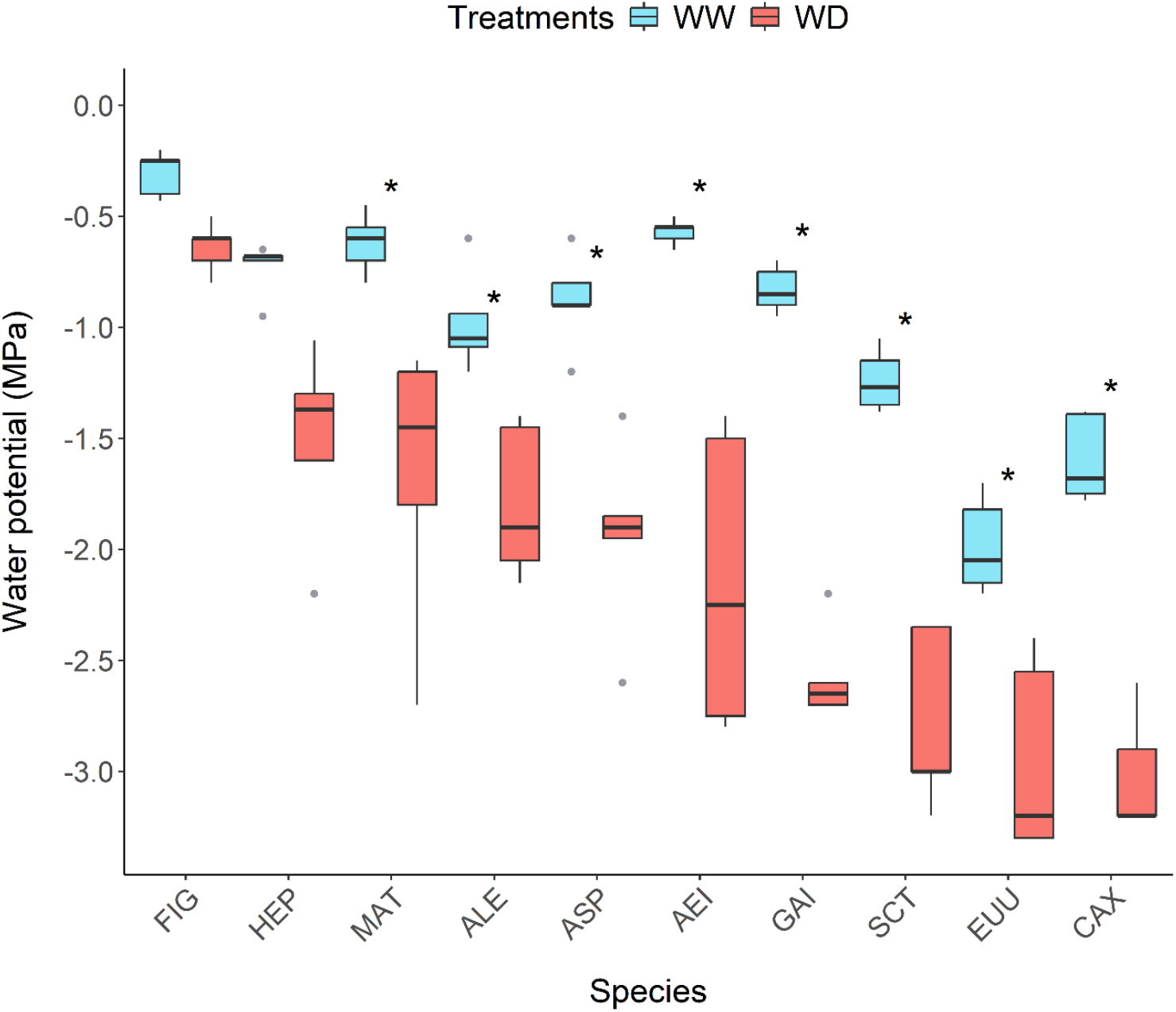
Water potential of seedlings from ten native species of the Semideciduous Seasonal Forest under well-watered (WW) and water-deficit (WD) conditions, measured at the point when stomatal conductance had decreased by ≥50% from its initial value in a severe WD experiment conducted in a greenhouse. AEI-*Aegiphila integrifolia* (Jacq.) Moldenke., ALE-*Allophylus edulis* (A.St.-Hil. *et al*.) Hieron. ex Niederl., (ALE), ASP-*Aspidosperma polyneuron* Müll. Arg., CAX-*Campomanesia xanthocarpa* (Mart.) O.Berg. EUU-*Eugenia uniflora* L., FIG-*Ficus guaranitica* Chodat, GAI -*Gallesia integrifolia* (Spreng.) Harms, HEP-*Heliocarpus popayanensis* Kunth, MAT-*Maclura tinctoria* (L.) D.Don ex Steud., SCT-*Schinus terebinthifolius* Raddi.

Spearman correlations using data from the final *g*_*s*_ measurement taken after a 50% reduction from initial *g*_s_ and the corresponding Ψw values indicated a moderate and significant correlation in the group under WW conditions (*rho* = 0.46, *p* < 0.01). In contrast, no significant correlation was found in the WD group (*rho* = 0.16, *p* = 0.26) (Fig. 4).

**Fig. 4.**
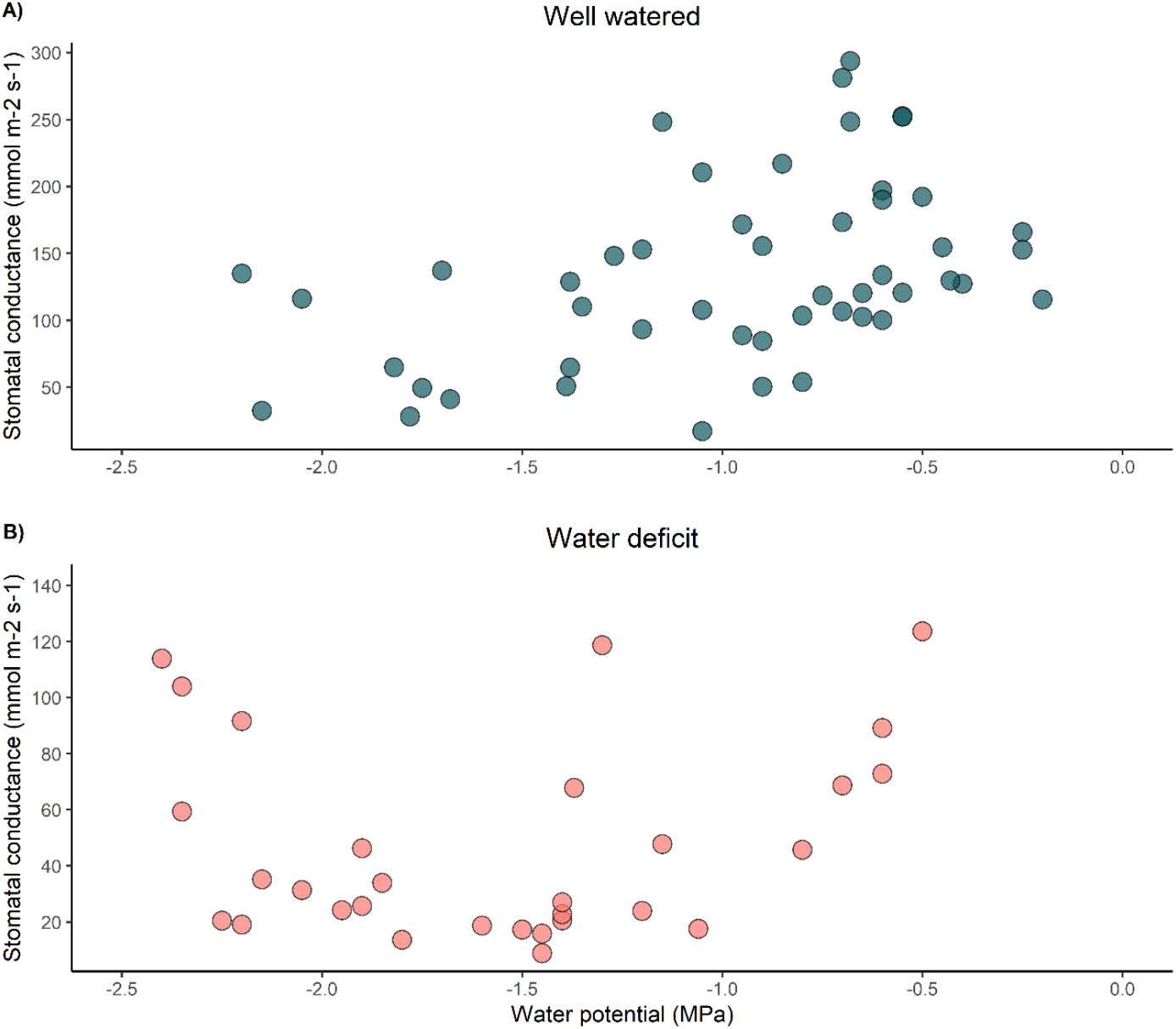
Spearman correlation between stem water potential and the final measurement of stomatal conductance (*g*_s_) (after a ≥50% reduction from initial *g*_s_) in individuals of 30 native species from the Semideciduous Seasonal Forest subjected to severe water deficit experiments in a greenhouse. A) Well watered conditions (*rho* = 0.46, *p* < 0.01). B) Water deficit conditions (*rho* = 0.16, *p* = 0.26).

## 4 DISCUSSION

In this study, we adapted a method standardized by Marchin et al. (2020). This type of experiment, in which a gradual and moderate WD is implemented on the plants, was introduced more than 50 years ago by Haan and Barfield (1971) and later modified by Fernández and Reynolds (2000). Here, we use “floral foam” (also used by Marchin et al. 2020 in their experiments) to apply, for the first time, a severe WD on nursery seedlings produced specifically for restoration projects. Those seedlings can be found easily in forest restoration-oriented commercial nurseries.

The great challenge for using these seedlings is to apply a severe WD on them just by ceasing irrigation, as is commonly done (Björkman and Powles, 1984; Souza et al., 2010), given that the water content in the 50 ml tube would be depleted in a matter of hours, making it impossible to reliably analyze ecophysiological responses of plants under stress. Another possibility would be to slowly reduce the irrigation (Do Carmo et al., 2021, Tiepo et al., 2018), but the amount of water poured into the tubes would need to be adjusted for each individual to keep all seedlings at the same wilting stage for a longer period, which would make it impossible to carry out these experiments with a large number of species in a short time. The present solution enables the study of many species simultaneously, thereby supporting predictive modeling of how remnant forests and restoration areas may respond to climate change.

The GLM results pointed out that the capillary irrigation method, with the use of floral foam blocks, was efficient in generating a controlled WD on the seedlings in the greenhouse, in maintaining the irrigation also controlled in the seedlings of the control group. Thus, it was possible to identify different levels of ecophysiological responses to WD among the species, which reflects different types of physiological strategies in water stress situations. Such interspecific variation provides insight into the relative drought resistance of different species.

In this regard, *F. guaranitica* and *C. xanthocarpa*, which used opposite strategies for water use, presented interesting results. While *F. guaranitica* reached 50% of the ig_s_ on the third day, *C. xanthocarpa* had a slower stomatal response, reaching 50% of the i*g*_s_ only on the fourth day of the experiment. The results of these strategies on Ψw in WD were also the opposite. *F. guaranitica* exhibited the highest Ψw in WD (averaging -0.64 MPa; Supplementary Material), being one of the few species that did not show a significant difference in Ψw between treatments, while *C. xanthocarpa* showed the lowest values (averaging -3.2 MPa; Supplementary Material). These results suggest that *F. guaranitica* behaved in an isohydric manner and *C. xanthocarpa* in an anisohydric manner, however, it is remarkable that the *g*_s_ of *F. guaranitica* even after a 50% reduction from its initial value, remained higher than the *g*_s_ of *C. xanthocarpa* under WW conditions, reflecting contrasting water use strategies.

Another noteworthy response was observed in *H. popayanensis*, which exhibited the highest *g*_*s*_ values under both WD and WW conditions on the third day of the experiment. This species reached a 50% reduction in g_s_ only on the fourth day and, similar to *F. guaranitica*, showed no significant difference in Ψw between treatments. These results indicate that, contrary to the suggestion by McDowell et al. (2008), no relationship between stomatal control and Ψw was identified in this study. This was further supported by the lack of correlation between the final *g*_s_ measurement − taken when *g*_s_ reached 50% or less of its initial value − and the corresponding Ψw in the WD group (*rho* = 0.16, p = 0.26). This result supports the findings of Martínez-Vilalta and Garcia-Forner (2017), who, in a review involving 33 species from Mediterranean and dry climates, concluded that strict regulation of Ψw is not necessarily associated with stronger stomatal control.

So, despite the classification proposed by McDowell et al. (2008), it seems to be more reasonable to use the terms iso/aniso-hydric to refer to the ability to keep Ψw relatively constant or not under climatic variations without suggesting a direct association with stomatal behavior (Martínez-Vilalta and Garcia-Forner, 2017). Thus, according to the criteria used in this study, the species that demonstrated greater resistance to WD were *H. popayanensis* and *F. guaranitica*. Contrastingly, *C. xanthocarpa, E. uniflora* were least resistant to WD, presenting very low Ψw when subjected to severe water stress. Meanwhile, most species were distributed along a gradient of responses to water deficit.

It is important to highlight that, alternative approaches to assessing drought resistance − such as measuring the time to mortality (Tyree et al., 2003, Poorter and Markesteijn, 2008, Slot and Poorter, 2007) − may be necessary to validate the findings presented here. Nevertheless, although Ψw cannot be used in isolation as a proxy for overall drought vulnerability (Martínez-Vilalta and Garcia-Forner, 2017), it remains one of the most reliable and widely used indicators for assessing plant resistance to WD. Additionally, other factors contributing to plant mortality under drought, such as carbon starvation and insect attack (McDowell et al., 2011), were not assessed in this study. Both carbon starvation and hydraulic failure depend on the intensity and duration of drought events (McDowell et al., 2008). Therefore, further studies considering varying levels and durations of WD are essential for a more comprehensive understanding of drought resistance in these species.

Moreover, it is important to note that plants exhibit a range of drought-related adaptations that were not examined in this study, including changes in leaf area (Varone et al., 2012), root exudate profiles (Brunn et al., 2022), osmotic adjustment (Ozturk et al., 2020, Yang et al. 2021), among others. Future research should focus on identifying functional traits and other physiological responses linked to drought resistance, which could enhance our understanding of this complex phenomenon.

Although further investigation is required, the approach adopted in this study enabled the rapid and relatively low-cost assessment of resistance to severe WD in ten native species of the SSF. With minor modifications, the method presented here can also be applied to induce a more moderate and prolonged water deficit in nursery seedlings, thus enabling the assessment of additional drought-related mechanisms, as demonstrated by Machin et al. (2020) using plants in 6 L pots. The possibility to conduct this type of experiment using nursery seedlings grown in 50 ml tubes, combined with the evaluation of responses that are relatively easy to measure, enables the simultaneous study of multiple species. In this way, the adapted method proposed in this study is expected to contribute to accelerating the acquisition of knowledge on how native species respond to water deficit.

## Supplementary Information

The present version contains supplementary material

## Acknowledgements

Authors are grateful to Henrique Garcia Rocha for providing the seedlings used in the experiments. We also thank the members of the Biodiversity and Ecosystem Restoration Laboratory and the Plant Ecophysiology Laboratory at the State University of Londrina for their assistance during the experiments.

## Author contributions

All authors agree to submission of the manuscript. LCDR and JMDT conceived the ideas and the overall study design; LCDR and JAP did the experimental work and collected the data; LCDR and FA performed statistical analysis; LCDR and JMDT wrote the first draft; all authors edited the manuscript, contributed critically to the drafts and gave final approval for publication.

## Data availability

The datasets generated and/or analyzed during the current study are available from the corresponding author upon request.

