## Additional figures and tables for "A method for estimating the response of nursery-grown Atlantic Forest tree seedlings to water deficit"

**Brazilian Journal of Botany.**

Larissa Cerqueira Dias Rodrigues[1], José Antônio Pimenta[1], Fátima Arcanjo[2], Alba Lúcia Cavalheiro[1], Halley Caixeta de Oliveira[1], José Marcelo Domingues Torezan[1*]

[1] Universidade Estadual de Londrina, Centro de Ciências Biológicas, Departamento de Biologia Animal e Vegetal– PR 445 Km 380– Campus Universitário, 86057-970, Londrina, PR, Brasil.

[2] Instituto de Biologia, Universidade Federal de Uberlândia, Rua Ceará, 1720, 38405-302, Uberlândia, MG, Brasil.

**Fig. S1** Plastic tube used for native tree species seedling production and “floral foam” blocks used to assembly the water deficit experiment. Note the wetness of the tube recently removed from the well in the foam block and the water in the bottom of the well.

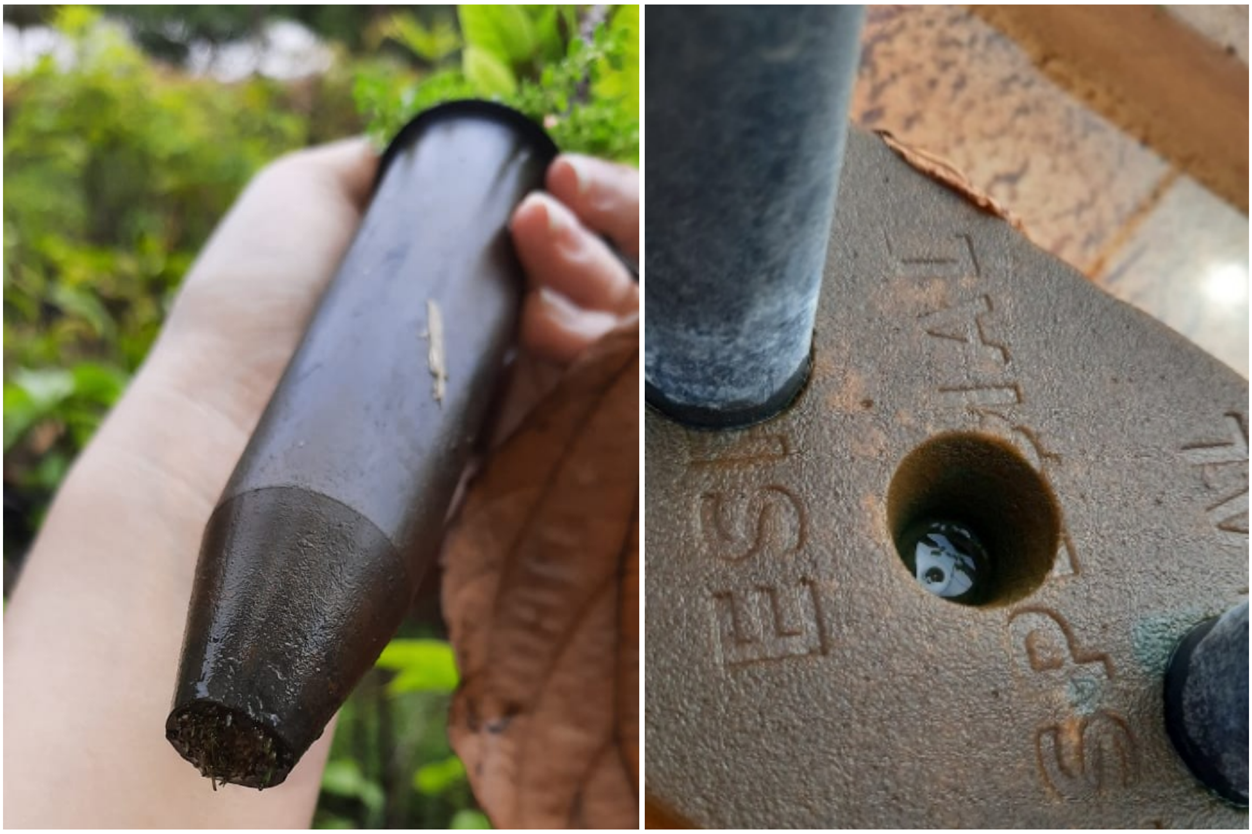

**Fig. S2** Method for allocating wells in floral foams, for the experiment of water deficit in a greenhouse. The seedlings will be inserted into the floral foam wells, which act as a solid column of low water permeability inside a plastic container filled with water; capillary irrigation is used to control the water content of the soil in the tubes with the seedlings. A) 2 cm distance from the edge to make the wells; B) marking on the tube used to make the wells, aiming to standardize their depth; C) wells placed in an interspersed pattern to avoid overlapping the tops of the seedlings used in the experiment. The red arrow indicates the marking made at the base of the tube, which delimited the depth of the wells.

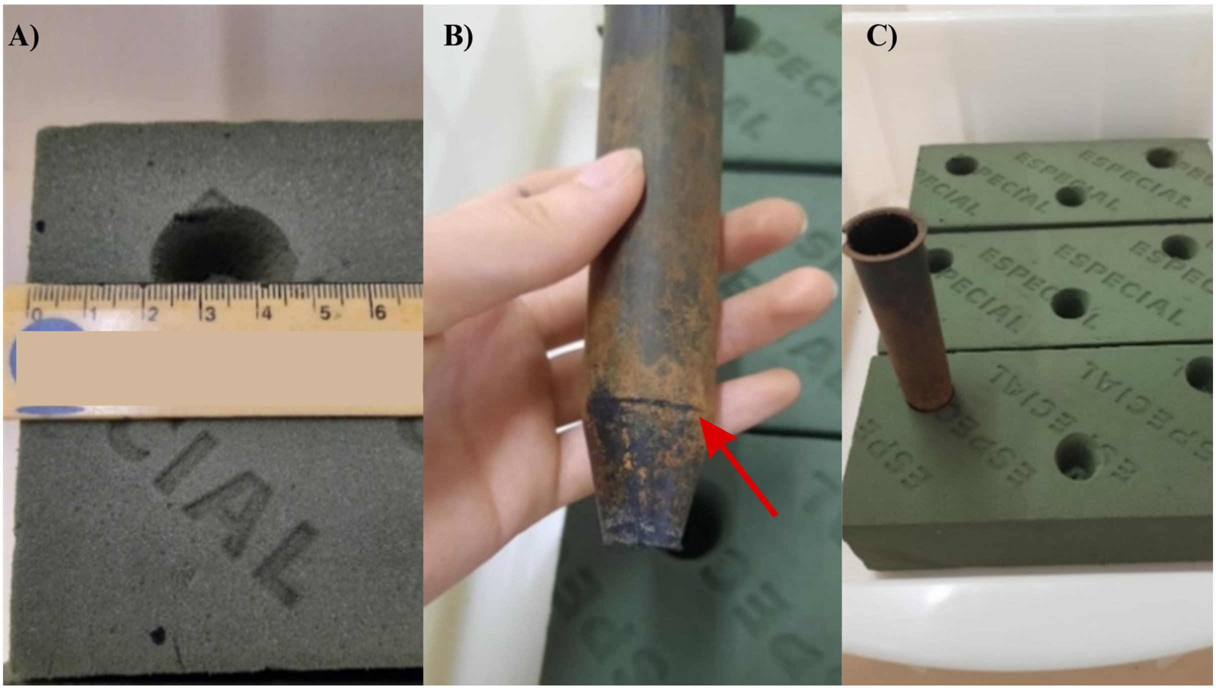

**Table S1** Comparison between Generalized Linear Models (GLMs) and Generalized Linear Mixed Models (GLMMs). Df: Degrees of freedom; AIC: Akaike Information Criterion; BIC: Bayesian Information Criterion; logLik: Log-likelihood; Dev: Model deviance; $\chi^2$: Chi-squared; $\chi^2$ Df: Degrees of freedom for the Chi-squared test; p($\chi^2$): p-value of the test. $g_{s}$: stomatal conductance; Ψw: water potential.

| **Models** | **Df** | **AIC** | **BIC** | **logLik** | **dev** | **X²** | **X² Df** | **p(χ2):** |
|---|---|---|---|---|---|---|---|---|
| GLM: $g_s$ ~ species * treatments | 21 | 1018.1 | 1072.6 | -488.07 | 976.15 | | | |
| GLM: $g_s$ ~ species * treatments + (1 \| experiment) | 22 | 1022.1 | 1079.2 | -489.07 | 978.15 | 0 | 1 | 1 |
| GLM: *log* Ψw ~ species * treatments | 17 | -62.97 | -24.75 | 48.48 | -96.97 | | | |
| GLM: *log* Ψw ~ species * treatments + (1 \| experiment) | 18 | -58.97 | -18.55 | 47.48 | -94.97 | 0 | 1 | 1 |

**Table S2** Data on stomatal conductance ($g_s$) from the third day of the experiment, the final $g_s$ measurement, and stem water potential (Ψw) after a ≥ 50% drop in initial $g_s$ for 10 native species of the Seasonal Semideciduous Forest subjected to severe water deficit experiments in a greenhouse (*n*= 5 individuals per species per traitment). Treat.: Treatments; WW – well watered individuals; WD – individuals subjected to water deficit.; Sd: standard deviation.

| **Species** | **Treat** | **$g_s$ (third day)** | | **Last $g_s$** | | **Ψw** | |
|---|---|---|---|---|---|---|---|
| | | **Mean** | **Sd** | **Mean** | **Sd** | **Mean** | **Sd** |
| *Aegiphila integrifolia* (Jacq.) Moldenke | WW | 158.00 | 31.45 | 173.14 | 61.14 | -0.57 | 0.06 |
| *Allophylus edulis* (A.St.-Hil. *et al.*) Hieron. ex Niederl | WW | 90.52 | 48.80 | 99.26 | 52.70 | -0.98 | 0.26 |
| *Aspidosperma polyneuron* Müll. Arg. | WW | 72.50 | 31.57 | 76.65 | 22.87 | -0.88 | 0.22 |
| *Campomanesia xanthocarpa* (Mart.) O.Berg | WW | 42.64 | 13.43 | 47.04 | 13.41 | -1.60 | 0.20 |
| *Eugenia uniflora L.* | WW | 86.72 | 34.09 | 97.24 | 46.34 | -1.98 | 0.22 |
| *Ficus guaranitica* Chodat | WW | 138.38 | 20.45 | 138.38 | 20.45 | -0.31 | 0.10 |
| *Gallesia integrifolia* (Spreng.) Harms | WW | 176.46 | 62.69 | 137.50 | 50.70 | -0.83 | 0.10 |
| *Heliocarpus popayanensis* Kunth | WW | 216.00 | 57.56 | 223.10 | 74.41 | -0.73 | 0.12 |
| *Maclura tinctoria* (L.) D.Don ex Steud. | WW | 165.84 | 42.00 | 174.86 | 54.04 | -0.62 | 0.14 |
| *Schinus therebentifolius* Raddi | WW | 169.27 | 58.04 | 169.27 | 58.04 | -1.24 | 0.14 |
| *Aegiphila integrifolia* | WD | 61.20 | 22.57 | 21.58 | 4.14 | -2.14 | 0.67 |
| *Allophylus edulis* | WD | 51.22 | 25.94 | 26.20 | 7.52 | -1.79 | 0.35 |
| *Aspidosperma polyneuron* | WD | 62.30 | 9.85 | 33.04 | 8.52 | -1.94 | 0.43 |
| *Campomanesia xanthocarpa* | WD | 35.80 | 15.39 | 44.52 | 28.84 | -3.02 | 0.27 |
| *Eugenia uniflora* | WD | 73.86 | 60.94 | 46.72 | 40.61 | -2.95 | 0.44 |
| *Ficus guaranitica* | WD | 80.06 | 28.91 | 80.06 | 28.91 | -0.64 | 0.11 |
| *Gallesia integrifolia* | WD | 53.23 | 38.48 | 19.13 | 4.12 | -2.57 | 0.21 |
| *Heliocarpus popayanensis* | WD | 148.66 | 47.39 | 62.90 | 44.65 | -1.51 | 0.43 |
| *Maclura tinctoria* | WD | 48.52 | 25.80 | 22.20 | 15.32 | -1.66 | 0.64 |
| *Schinus therebentifolius* | WD | 74.49 | 48.06 | 74.49 | 48.06 | -2.78 | 0.40 |

**Table S3** Results of the GLM analysis of stomatal conductance $g_s$ data from 10 native species of the Semideciduous Seasonal Forest, measured on the third day of severe drought stress in a greenhouse. The effects of the explanatory variables – species and treatments (well-watered and water deficit, WD) – on $g_s$ were estimated using a Gamma distribution.

| **Response Variable** | **Explanatory Variable** | **Estimate** | **SE** | **t value** | **Pr(>\|t\|)** |
|---|---|---|---|---|---|
| $g_s$ of third day of experiment | *Aegiphila integrifolia* **(Intercept)** | 0.01 | 0.00 | 5.11 | 0.00* |
| | *Allophylus edulis* | 0.00 | 0.00 | 1.89 | 0.06 |
| | *Aspidosperma polyneuron* | 0.01 | 0.00 | 2.51 | 0.01* |
| | *Campomanesia xanthocarpa* | 0.02 | 0.00 | 3.60 | 0.00* |
| | *Eugenia uniflora* | 0.01 | 0.00 | 2.02 | 0.05* |
| | *Ficus guaranitica* | 0.00 | 0.00 | 0.48 | 0.63 |
| | *Gallesia integrifolia* | 0.00 | 0.00 | -0.40 | 0.69 |
| | *Heliocarpus popayanensis* | 0.00 | 0.00 | -1.11 | 0.27 |
| | *Maclura tinctoria.* | 0.00 | 0.00 | -0.18 | 0.86 |
| | *Schinus terebinthifolia* | 0.00 | 0.00 | -0.25 | 0.80 |
| | WD | 0.01 | 0.00 | 2.92 | 0.00* |
| | *Aegiphila sellowiana*: WD | 0.00 | 0.01 | -0.28 | 0.78 |
| | *Allophylus edulis*: WD | -0.01 | 0.01 | -1.44 | 0.15 |
| | *Aspidosperma polyneuron:* WD | -0.01 | 0.01 | -0.70 | 0.49 |
| | *Campomanesia xanthocarpa:* WD | -0.01 | 0.00 | -1.64 | 0.11 |
| | *Eugenia uniflora:* WD | 0.00 | 0.00 | -1.07 | 0.29 |
| | *Ficus guaranitica:* WD | 0.00 | 0.01 | 0.57 | 0.57 |
| | *Gallesia integrifolia:* WD | -0.01 | 0.00 | -2.09 | 0.04* |
| | *Heliocarpus popayanensis:* WD | 0.00 | 0.01 | 0.84 | 0.40 |
| | *Maclura tinctoria:* WD | 0.00 | 0.00 | -0.56 | 0.58 |

Note: * Indicates significant differences.

**Table S4** Post-hoc multiple comparisons of the stomatal conductance on the third day of experiments among water deficit individuals across species. Emmean – Estimated marginal mean, the adjusted mean response while controlling for other factors in the model; SE – Standard error; Df – Degrees of freedom; Asymp LCL – Lower bound of the asymptotic confidence interval; Asymp UCL – Upper bound of the asymptotic confidence interval; Group – Species sharing the same letters are statistically similar.

| **Species** | **Emmean** | **SE** | **Df** | **Asymp. LCL** | **Asymp. UCL** | **Groups** |
|---|---|---|---|---|---|---|
| *Heliocarpus popayanensis* | 0.01 | 0.00 | 79 | 0.00 | 0.01 | a |
| *Ficus guaranitica* | 0.01 | 0.00 | 79 | 0.01 | 0.02 | ab |
| *Schinus terebinthifolius* | 0.01 | 0.00 | 79 | 0.01 | 0.02 | ab |
| *Eugenia uniflora* | 0.01 | 0.00 | 79 | 0.01 | 0.02 | ab |
| *Aspidosperma polyneuron* | 0.02 | 0.00 | 79 | 0.01 | 0.03 | ab |
| *Aegiphila integrifolia* | 0.02 | 0.00 | 79 | 0.01 | 0.03 | ab |
| *Gallesia integrifolia* | 0.02 | 0.00 | 79 | 0.01 | 0.03 | ab |
| *Allophylus edulis* | 0.02 | 0.00 | 79 | 0.01 | 0.03 | ab |
| *Maclura tinctoria* | 0.02 | 0.00 | 79 | 0.01 | 0.03 | ab |
| *Campomanesia xanthocarpa* | 0.03 | 0.01 | 79 | 0.01 | 0.04 | b |

**Table S5** Post-hoc multiple comparisons of the stomatal conductance on the third day of experiments for each species across treatments. Treat – treatment, WW – well watered, WD – water deficit, Emmean – estimated marginal mean, adjusted mean response controlling for other factors in the model; SE – standard error; Df – degrees of freedom; Asymp LCL – lower bound of the asymptotic confidence interval; Asymp UCL – upper bound of the asymptotic confidence interval; Group – treatments sharing the same letters are statistically similar.

| **Species** | **Treat** | **Emmean** | **SE** | **Gl** | **Asymp LCL** | **Asymp UCL** | **Grupos** |
|---|---|---|---|---|---|---|---|
| *Aegiphila integrifolia* | WW | 0.01 | 0.00 | 79 | 0.00 | 0.01 | a |
| | WD | 0.02 | 0.00 | 79 | 0.01 | 0.02 | b |
| *Allophylus edulis* | WW | 0.01 | 0.00 | 79 | 0.01 | 0.02 | a |
| | WD | 0.02 | 0.00 | 79 | 0.01 | 0.03 | a |
| *Aspidosperma polyneuron* | WW | 0.01 | 0.00 | 79 | 0.01 | 0.02 | a |
| | WD | 0.02 | 0.00 | 79 | 0.01 | 0.02 | a |
| *Campomanesia xanthocarpa* | WW | 0.02 | 0.00 | 79 | 0.01 | 0.03 | a |
| | WD | 0.03 | 0.01 | 79 | 0.02 | 0.04 | a |
| *Eugenia uniflora* | WW | 0.01 | 0.00 | 79 | 0.01 | 0.02 | a |
| | WD | 0.01 | 0.00 | 79 | 0.01 | 0.02 | a |
| *Ficus guaranitica* | WW | 0.01 | 0.00 | 79 | 0.00 | 0.01 | a |
| | WD | 0.01 | 0.00 | 79 | 0.01 | 0.02 | a |
| *Gallesia integrifolia* | WW | 0.01 | 0.00 | 79 | 0.00 | 0.01 | a |
| | WD | 0.02 | 0.00 | 79 | 0.01 | 0.03 | b |
| *Heliocarpus popayanensis* | WW | 0.00 | 0.00 | 79 | 0.00 | 0.01 | a |
| | WD | 0.01 | 0.00 | 79 | 0.00 | 0.01 | a |
| *Maclura tinctoria* | WW | 0.01 | 0.00 | 79 | 0.00 | 0.01 | a |
| | WD | 0.02 | 0.00 | 79 | 0.01 | 0.03 | b |
| *Schinus therebentifolius* | CC | 0.01 | 0.00 | 79 | 0.00 | 0.01 | a |
| | DH | 0.01 | 0.00 | 79 | 0.01 | 0.02 | b |

**Table S6** GLM results of stem water potential (Ψw) in 10 native species of the Semideciduous Seasonal Forest, subjected to severe water deficit (WD) in a greenhouse. Analyses were performed using data from individuals that exhibited ≥50% loss in stomatal conductance. The effects of the explanatory variables – species and treatment (well-watered and WD) – on Ψw were estimated using a GLM with Gamma distribution.

| **Response Variable** | **Explanatory Variable** | **Estimate** | **Std. Error** | **t value** | **Pr(>\|t\|)** |
|---|---|---|---|---|---|
| *log* Ψw | *Aegiphila integrifolia* **(Intercept)** | 0.97 | 0.06 | 17.61 | < 0.01 |
| | *Allophylus edulis* | -0.11 | 0.07 | -1.51 | 0.14 |
| | *Aspidosperma polyneuron* | -0.07 | 0.08 | -0.94 | 0.35 |
| | *Campomanesia xanthocarpa* | 0.51 | 0.10 | 5.05 | 0.00* |
| | *Eugenia uniflora* | 0.46 | 0.10 | 4.65 | 0.00* |
| | *Ficus guaranitica* | -0.29 | 0.07 | -4.35 | 0.00* |
| | *Gallesia integrifolia* | 0.16 | 0.08 | 1.87 | 0.07 |
| | *Heliocarpus popayanensis* | -0.17 | 0.07 | -2.35 | 0.02* |
| | *Maclura tinctoria .* | -0.13 | 0.07 | -1.80 | 0.08 |
| | *Schinus terebinthifolia* | 0.30 | 0.09 | 3.32 | 0.00* |
| | WD | -0.30 | 0.07 | -4.47 | 0.00* |
| | *Aegiphila integrifolia*: WD | 0.16 | 0.09 | 1.67 | 0.10 |
| | *Allophylus edulis*: WD | 0.11 | 0.09 | 1.13 | 0.26 |
| | *Aspidosperma polyneuron:* WD | -0.36 | 0.12 | -3.09 | 0.00* |
| | *Campomanesia xanthocarpa:* WD | -0.22 | 0.12 | -1.88 | 0.06 |
| | *Eugenia uniflora:* WD | 0.27 | 0.09 | 3.12 | 0.00* |
| | *Ficus guaranitica:* WD | -0.13 | 0.10 | -1.28 | 0.20 |
| | *Gallesia integrifolia:* WD | 0.19 | 0.09 | 2.06 | 0.04* |
| | *Heliocarpus popayanensis:* WD | 0.14 | 0.09 | 1.50 | 0.14 |
| | *Maclura tinctoria:* WD | -0.22 | 0.11 | -2.04 | 0.05* |

Note: * Indicates significant differences.

**Table S7** Post-hoc multiple comparisons of stem water potential among water deficit individuals across species. Emmean – Estimated marginal mean, the adjusted mean response while controlling for other factors in the model; SE – Standard error; Df – Degrees of freedom; Asymp LCL – Lower bound of the asymptotic confidence interval; Asymp UCL – Upper bound of the asymptotic confidence interval; Group – Species sharing the same letters are statistically similar.

| **Species** | **Emmean** | **SE** | **Df** | **Asymp. LCL** | **Asymp. UCL** | **Groups** |
|---|---|---|---|---|---|---|
| *Ficus guaranitica* | 0.68 | 0.04 | 79 | 0.57 | 0.79 | a |
| *Heliocarpus popayanensis* | 0.80 | 0.05 | 79 | 0.67 | 0.94 | ab |
| *Maclura tinctoria* | 0.84 | 0.05 | 79 | 0.70 | 0.98 | ab |
| *Allophylus edulis* | 0.86 | 0.05 | 79 | 0.72 | 1.00 | abc |
| *Aspidosperma polyneuron* | 0.90 | 0.05 | 79 | 0.75 | 1.05 | bc |
| *Aegiphila integrifolia* | 0.97 | 0.06 | 79 | 0.81 | 1.13 | bcd |
| *Gallesia integrifolia* | 1.13 | 0.06 | 79 | 0.95 | 1.32 | cde |
| *Schinus terebinthifolius* | 1.27 | 0.07 | 79 | 1.07 | 1.48 | de |
| *Eugenia uniflora* | 1.43 | 0.08 | 79 | 1.20 | 1.66 | e |
| *Campomanesia xanthocarpa* | 1.48 | 0.08 | 79 | 1.24 | 1.72 | e |

**Table S8** Post-hoc multiple comparisons of the stem water potential for each species across treatments Treat – treatment, WW – well watered, WD – water deficit, Emmean – estimated marginal mean, adjusted mean response controlling for other factors in the model; SE – standard error; Df – degrees of freedom; Asymp LCL – lower bound of the asymptotic confidence interval; Asymp UCL – upper bound of the asymptotic confidence interval; Group – treatments sharing the same letters are statistically similar.

| **Species** | **Treat** | **Emmean** | **SE** | **Gl** | **Asymp LCL** | **Asymp UCL** | **Grupos** |
|---|---|---|---|---|---|---|---|
| *Aegiphila integrifolia* | WW | 0.67 | 0.04 | 79 | 0.59 | 0.76 | a |
| | WD | 0.97 | 0.06 | 79 | 0.85 | 1.10 | b |
| *Allophylus edulis* | WW | 0.72 | 0.05 | 79 | 0.62 | 0.82 | a |
| | WD | 0.86 | 0.05 | 79 | 0.75 | 0.97 | b |
| *Aspidosperma polyneuron* | WW | 0.71 | 0.04 | 79 | 0.62 | 0.80 | a |
| | WD | 0.90 | 0.05 | 79 | 0.78 | 1.02 | b |
| *Campomanesia xanthocarpa* | WW | 0.82 | 0.05 | 79 | 0.71 | 0.92 | a |
| | WD | 1.48 | 0.08 | 79 | 1.29 | 1.67 | b |
| *Eugenia uniflora* | WW | 0.91 | 0.05 | 79 | 0.79 | 1.03 | a |
| | WD | 1.43 | 0.08 | 79 | 1.24 | 1.61 | b |
| *Ficus guaranitica* | WW | 0.65 | 0.04 | 79 | 0.56 | 0.73 | a |
| | WD | 0.68 | 0.04 | 79 | 0.59 | 0.77 | a |
| *Gallesia integrifolia* | WW | 0.70 | 0.04 | 79 | 0.61 | 0.79 | a |
| | WD | 1.13 | 0.06 | 79 | 0.98 | 1.28 | b |
| *Heliocarpus popayanensis* | WW | 0.69 | 0.04 | 79 | 0.60 | 0.78 | a |
| | WD | 0.80 | 0.05 | 79 | 0.70 | 0.91 | a |
| *Maclura tinctoria* | WW | 0.68 | 0.04 | 79 | 0.59 | 0.77 | a |
| | WD | 0.84 | 0.05 | 79 | 0.73 | 0.95 | b |
| *Schinus therebentifolius* | CC | 0.76 | 0.04 | 79 | 0.66 | 0.85 | a |
| | DH | 1.27 | 0.07 | 79 | 1.11 | 1.44 | b |
